# Stacks off tracks: A role for the golgin AtCASP in plant endoplasmic reticulum – Golgi apparatus tethering

**DOI:** 10.1101/115840

**Authors:** Anne Osterrieder, Stan W Botchway, Andy Ward, Tijs Ketelaar, Norbert de Ruijter, Chris Hawes

## Abstract

The plant Golgi apparatus modifies and sorts incoming proteins from the endoplasmic reticulum (ER), and synthesises cell wall matrix material. Plant cells possess numerous motile Golgi bodies, which are connected to the ER by yet to be identified tethering factors. Previous studies indicated a role of cis-Golgi plant golgins (long coiled-coil domains proteins anchored to Golgi membranes) in Golgi biogenesis. Here we show a tethering role for the golgin AtCASP at the ER-Golgi interface. Using live-cell imaging, Golgi body dynamics were compared in Arabidopsis thaliana leaf epidermal cells expressing fluorescently tagged AtCASP, a truncated AtCASP-ΔCC lacking the coiled-coil domains, and the Golgi marker STtmd. Golgi body speed and displacement were significantly reduced in AtCASP-ΔCC lines. Using a dual-colour optical trapping system and a TIRF-tweezer system, individual Golgi bodies were captured in planta. Golgi bodies in AtCASP-ΔCC lines were easier to trap, and the ER-Golgi connection was more easily disrupted. Occasionally, the ER tubule followed a trapped Golgi body with a gap, indicating the presence of other tethering factors. Our work confirms that the intimate ER-Golgi association can be disrupted or weakened by expression of truncated AtCASP-ΔCC, and suggests that this connection is most likely maintained by a golgin-mediated tethering complex.

**Highlight:** Here we show that the Golgi-associated *Arabidopsis thaliana* protein AtCASP may form part of a golgin-mediated tethering complex involved in anchoring plant Golgi stacks to the endoplasmic reticulum (ER).

## Introduction

The architecture of the Golgi apparatus is distinct and seemingly simple: An organelle composed of lipids and proteins, arranged as a polarised stack of flattened cisternae, capable of processing and distributing secretory cargo around and out of the cell (Klumperman, 2011; Polishchuk and Mironov, 2004; Staehelin and Moore, 1995). Yet, the exact mechanisms of Golgi stack assembly and maintenance are still not fully understood (Wang and Seemann, 2011). It is clear though that these processes depend on a highly complex and tightly regulated cascade of molecular events (Altan-Bonnet *et al.,* 2004), in which proteins attach to correct membranes and precisely orchestrate a multitude of tethering, fusing and budding events (Hawes *et al.,* 2010; Wilson and Ragnini-Wilson, 2010).

A Golgi stack has a *cis*-face through which it receives secretory cargo proteins from the endoplasmic reticulum (ER, Hawes *et al.,* 2008; Lorente-Rodriguez and Barlowe, 2011; Robinson *et al.,* 2015; Robinson *et al.,* 2007), and a *trans-*face where protein cargo exits via the trans-Golgi network and enters intracellular or exocytotic post-Golgi transport routes (Foresti and Denecke, 2008; Park and Jürgens, 2011).

Secretory cargo proteins move through the stack to be processed sequentially and glycosylated by residential N-glycosyltransferases (Schoberer and Strasser, 2011). COPII-coated membrane carriers function in anterograde ER-to-Golgi transport, whereas COPI-coated vesicles transport proteins backwards within the stack and from the *cis*-Golgi stack back to the ER for recycling (Robinson *et al.,* 2015).

To make matters more complicated, Golgi structure differs significantly between kingdoms. The mammalian Golgi apparatus is most often organised as a stationary perinuclear ‘Golgi ribbon’ in which single stacks appear to laterally fuse to create a ribbon-like structure (Nakamura *et al.,* 2012). Plant cells on the other hand contain numerous discrete and highly mobile Golgi bodies (Hawes and Satiat-Jeunemaitre, 2005), which move along the actin cytoskeleton (Boevink *et al*., 1998; Nebenführ *et al*., 1999) in a myosin-dependent manner (Sparkes, 2010).

In leaf epidermal cells, Golgi bodies and ER exit sites (specialised subdomains of the ER at which protein export occurs) appear intimately associated, resulting in the adoption of the “mobile secretory unit concept” (da Silva *et al.,* 2004; Hanton *et al.,* 2009; Robinson *et al.,* 2015). A study using optical tweezers in living leaf epidermal cells confirmed this concept by demonstrating a strong physical connection between ER tubules and Golgi bodies upon micromanipulation of the latter (Sparkes *et al.*, 2009b).

However, to date we have no definite information on the nature of the molecular complexes that are assumed to be involved in tethering Golgi stacks to ER exit sites. In mammalian cells the golgins, a family of Golgi-localised proteins with long coiled-coil domains, participate in tethering events at the Golgi (Barinaga-Rementeria Ramirez and Lowe, 2009; Barr and Short, 2003; Short *et al.,* 2005; Wong and Munro, 2014). Their coiled-coil domains form a rod-like structure that protrudes into the cytoplasm and thus are free to interact with membranous structures such as cargo carriers and neighbouring cisternae, or form a part of larger protein tethering complexes (Chia and Gleeson, 2014; Gillingham and Munro, 2003; Malsam and Söllner, 2011).

Plants possess a set of putative golgins that locate to Golgi bodies, and protein interaction partners have been identified for some of them (Gilson *et al.,* 2004; Latijnhouwers *et al.,* 2007; Latijnhouwers *et al.,* 2005b; Matheson *et al.,* 2007; Osterrieder, 2012; Renna *et al.,* 2005). Their subcellular functions largely remain unclear, although a mammalian p115 homologue has been suggested to be a tethering factor involved in anterograde transport from the ER (Takahashi *et al.,* 2010). A *cis*-Golgi localised golgin and good candidate protein for tethering Golgi bodies to ER exit sites is AtCASP (Latijnhouwers *et al.,* 2007; Latijnhouwers *et al.,* 2005a; Renna *et al.,* 2005), a type II transmembrane domain protein with a topology similar to the animal CASP protein (Gillingham *et al.,* 2002). Its N-terminal coiled-coil domains are predicted to form a rod-like structure reaching into the cytoplasm, whereas its C-terminus contains a transmembrane domain sufficient for Golgi targeting (Renna *et al.,* 2005) and multiple di-acidic DXE motifs required for ER export (Hanton *et al.,* 2005).

CASP, initially identified as a nuclear alternative splicing product of the CUTL1 gene encoding the transcriptional repressor CCAAT displacement protein CDP/cut (Lievens *et al.,* 1997), was found to locate to Golgi membranes by Gillingham and co-workers (2002). The authors observed protein interactions between CASP and golgin-84 and hSec23 at substochiometric levels, as well as genetic interactions between the yeast CASP homologue COY1 and the SNAREs Gos1p and Sec22p, suggesting a role for CASP in membrane trafficking. Subsequently, Malsam and colleagues reported CASP to function in an asymmetric tethering complex with Golgin-84, with CASP decorating Golgi membranes and Golgin-84 COPI vesicles (Malsam *et al.,* 2005).

Our previous studies indicated a role for AtCASP in Golgi stack formation at an early stage and possibly at the level of ER exit sites (Osterrieder *et al.,* 2010). After Golgi membrane disruption using an inducible GTP-locked version of the small COPII GTPase SAR1, GFP-AtCASP co-located with Sar1-GTP-YFP in punctate structures on the ER (Osterrieder *et al.,* 2010). AtCASP also labelled reforming Golgi bodies before Golgi membrane markers after washout of the secretory inhibitor Brefeldin A (Schoberer *et al.,* 2010).

In this study we used full-length and coiled-coil deletion mutant versions of AtCASP in conjunction with laser tweezers (Sparkes, 2016) to assess its potential role in ER-Golgi tethering and protein transport. Our findings implicate a role for AtCASP in tethering at the ER-Golgi interface, as over-expression of a dominant-negative truncation interferes with the stability of the ER-Golgi connection. However, our observations also suggest the involvement of additional and as yet uncharacterised tethering factors.

## Materials and Methods

### Molecular biology

Standard molecular techniques were used as described in Sambrook and Russel (2001). Fluorescent mRFP fusions of full-length AtCASP and truncated AtCASP-ΔCC were created using the previously published pENTR1A clones (Latijnhouwers *et al.,* 2007) and Gateway^®^ cloning technology according to instructions of the manufacturer (Life Technologies) into the binary expression vector pB7WGR2 (Karimi *et al.,* 2002). Constructs were sequenced and transformed into the *Agrobacterium tumefaciens* strain GV3101::mp90

### Transient expression of fluorescent protein fusions in tobacco plants

Transient expression of fluorescent protein fusions in tobacco leaves was carried out using *Agrobacterium*-mediated infiltration of *Nicotiana tabacum sp.* lower leaf epidermal cells (Sparkes *et al.,* 2006). Plants were grown in the greenhouse at 21 °C, and were used for *Agrobacterium tumefaciens* infiltration at the age of 5-6 weeks. Leaf samples were analysed 2-4 days after infiltration.

### Stable expression of Arabidopsis thaliana plants

Stable *Arabidopsis* plants were created using the *Agrobacterium-mediated* floral dip method (Clough and Bent, 1998). *Arabidopsis* plants from a stable GFP-HDEL line (Zheng *et al.,* 2004) were transformed either with mRFP-AtCASP or mRFP-AtCASP-ΔCC and grown on solid ½ MS medium with BASTA selection. All experiments were performed in T3 or T4 seedlings. As control, the previously described *Arabidopsis* line expressing the Golgi marker STtmd-mRFP and the ER marker GFP-HDEL was used (Sparkes *et al.,* 2009b).

### Confocal laser scanning microscopy

High-resolution confocal images were obtained using an inverted Zeiss LSM510 Meta confocal laser scanning microscope (CLSM) microscope and a 40x, 63x or 100x oil immersion objective. For imaging GFP in combination with mRFP, an Argon ion laser 488 nm and a HeNe ion laser 543 nm were used with line switching, using the multitrack facility of the CLSM. Fluorescence was detected using a 488/543 dichroic beam splitter, a 505-530 band pass filter for GFP and a 560-615 band pass filter for mRFP.

### Optical trapping

Optical trapping was carried out in stable *Arabidopsis* lines, using 1) a commercially available dual colour system at Wageningen University, The Netherlands, comprising a 1063nm, 3000mW Nd:YAG laser (Spectra Physics) and x-y galvo scanner (MMI, Glattbrugg, Switzerland) attached to a Zeiss Axiovert 200M with a Zeiss LSM510 Meta confocal laser scanning system (Sparkes *et al.,* 2009b), and 2) a custom-built TIRF-Tweezer system at the Central Laser Facility, Harwell (Gao *et al.,* 2016).

Golgi bodies were trapped using a 1090 nm infrared laser with intensity between 50 and 130 mW. For the ‘100 Golgi test’, Golgi bodies were scored as being trapped if they could be moved by the laser beam.

### Latrunculin B treatment

To inhibit actin-myosin based Golgi movement which was required during optical trapping confocal microscopy, *Arabidopsis* cotyledonary leaves were treated with the actin-depolymerising agent 2.5 μΜ latrunculin B for 30 min as previously described (Sparkes *et al.,* 2009b). Optical trapping experiments were performed within a time scale of two hours after latrunculin B application.

### Tracking and statistical analysis of Golgi body and ER dynamics

Movies for analysis of Golgi body dynamics in stable *Arabidopsis* lines were taken with a 63x PlanApo 1.4 NA oil objective at 512x512 resolution, optical zoom of 3.7 over a region of interest sized 244x242 pixels, and recorded for 50 frames at 0.9 frames/sec. Individual Golgi bodies were tracked using Fiji (Schindelin, *et al.* 2012) and the tracking plugin MTrackJ (Meijering, *et al.* 2012). Average Golgi body displacement and speed per cell were calculated from the median values using Microsoft Excel. Track lengths of trapped Golgi bodies and tracks in relation to tips of ER tubules were analysed using ImageJ (Schneider *et al.* 2012) and MTrackJ.

Statistical analysis of average displacement and speed in control and mutant cells were performed in Graphpad Prism through one-way ANOVA analysis followed by unpaired two-tailed type 2 Student t-tests. Statistical analysis of Golgi body trapping in control and mutant lines trapped within the ‘100 Golgi test’ was performed on numerical values in Microsoft Excel, using a Chi-Square test.

## Results

*Fluorescently labelled full-length and mutant AtCASP constructs co-locate with the Golgi marker STtmd-GFP in tobacco leaf epidermal cells*

The first step in assessing the function of a putative protein tether is to disturb its tethering capability. If AtCASP played a role in tethering events between the ER and Golgi bodies, deleting its coiled-coil domain could be predicted to affect Golgi morphology, function or dynamics, possibly resulting in changes in: 1) the subcellular location of the fluorescent mutant compared to the full-length protein, 2) the subcellular location of Golgi bodies in relation to the endoplasmic reticulum, 3) Golgi body dynamics, such as speed or displacement, or 4) in the physical interaction between Golgi bodies and the endoplasmic reticulum tested by optical tweezer based displacement of Golgi bodies.

The mRFP (monomeric red fluorescent protein) constructs used for this study were full-length AtCASP and a deletion mutant AtCASP-ΔCC. In this mutant, the coiled-coil domains were deleted to produce a truncated protein. AtCASP-ΔCC consists of the C-terminal 463 base pairs, including a transmembrane domain that confers Golgi localisation (Figure 1a, Latijnhouwers *et al.,* 2007; Renna *et al.,* 2005). This construct may act as a dominant-negative mutant, as it competes with endogenous wild type AtCASP for Golgi membrane insertion, but lacks the native protein’s potential tethering functions. As previously described for the green fluorescent protein (GFP) versions (Latijnhouwers *et al.,* 2007; Renna *et al.,* 2005), upon transient expression in tobacco leaf epidermal cells, the full-length mRFP-AtCASP co-located with the standard Golgi marker STtmd-GFP (Boevink *et al.,* 1998) in punctate structures (Figure 1b). Similarly, mRFP-AtCASP-ΔCC contained sufficient information to target the fluorescent fusion protein to Golgi bodies (Figure 1c). No obvious changes in localization were observed between the full-length and the mutant AtCASP construct.

**Figure 1.**
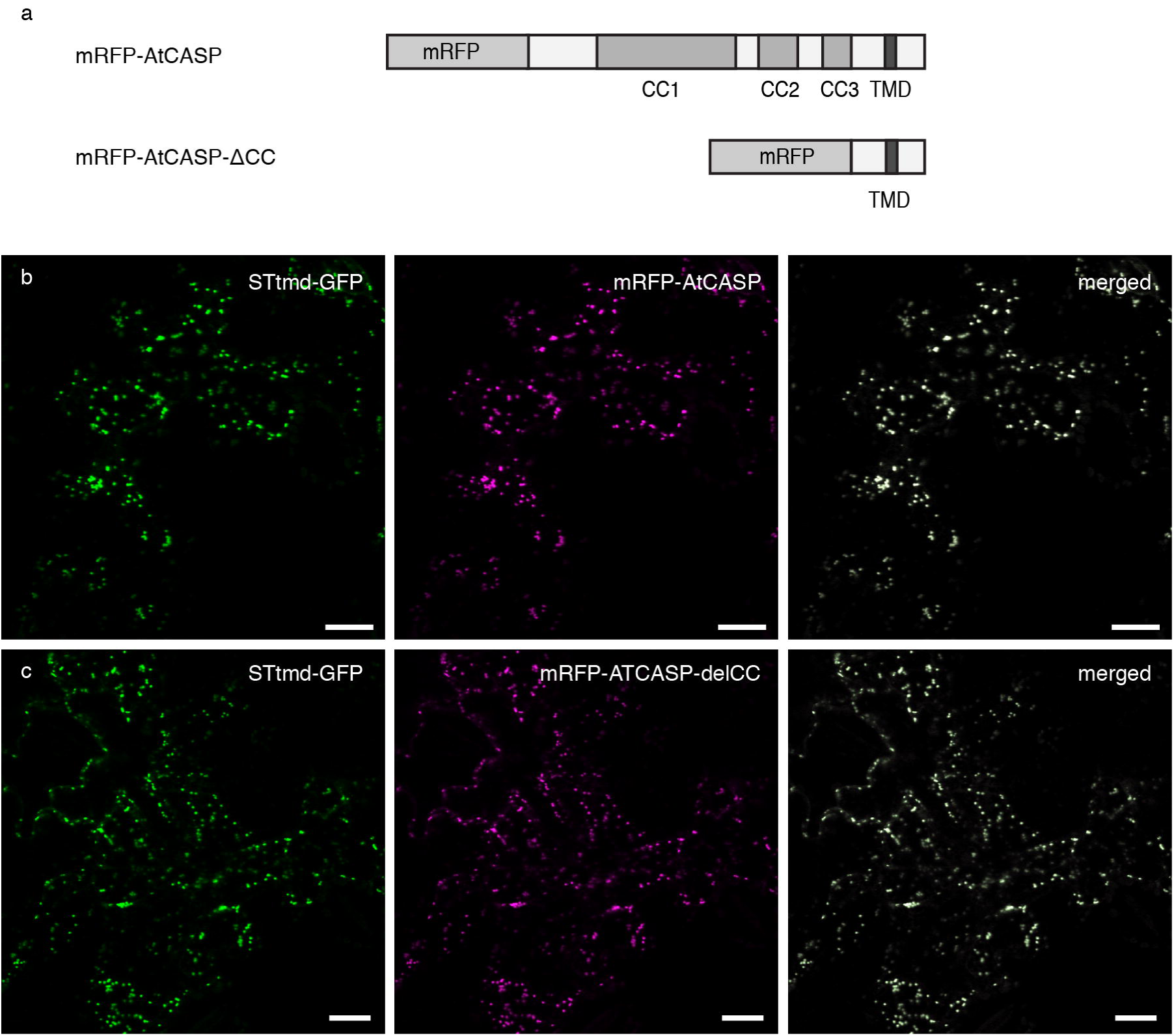
Fluorescent AtCASP full-length and mutant constructs. a) Diagram depicting the domain structure of fluorescent AtCASP constructs used in this study: full-length AtCASP and a truncation AtCASP-ΔCC consisting of its C-terminus (463 base pairs including the transmembrane domain), but missing the coiled-coil domains which convey its tethering function. CC = coiled-coil domain, TMD = transmembrane domain, mRFP = monomeric red fluorescent protein.
b) and c) Confocal laser scanning micrographs of tobacco leaf epidermal cells three days after transfection, transiently expressing the standard Golgi marker STtmd-GFP (green) and b) full-length mRFP-AtCASP or c) of mRFP-AtCASP-ΔCC (magenta). Both constructs co-locate in punctate structures which represent Golgi bodies. Cells were transfected using agrobacterium-mediated transformation. STtmd-GFP was infiltrated at OD600 = 0.05, mRFP-AtCASP constructs at OD_600_ = 0.1. Scale bars = 20 μm.

### Golgi body speed and displacement are significantly reduced in mutant AtCASP lines

To obtain qualitative and quantitative data on any changes in interactions between Golgi bodies and the ER, stable *Arabidopsis* lines expressing full-length mRFP-AtCASP or mRFP-AtCASP-ΔCC in a GFP-HDEL background were created. A transgenic *Arabidopsis* line expressing the Golgi marker STtmd-mRFP and the ER marker GFP-HDEL (Runions *et al.,* 2006; Sparkes *et al.,* 2009b) was used as a control (Figure 2a). Cotyledonary leaf epidermal cells in 4-6 day old seedlings were analysed using confocal laser scanning microscopy. Neither the mRFP-AtCASP/GFP-HDEL lines (Figure 2b) nor the mutant mRFP-AtCASP-ΔCC/GFP-HDEL lines (Figure 2c) displayed any obvious differences, at the resolution of the confocal microscope, in Golgi morphology, or spatial positioning relative to the surface of the ER, compared to the control.

**Figure 2.**
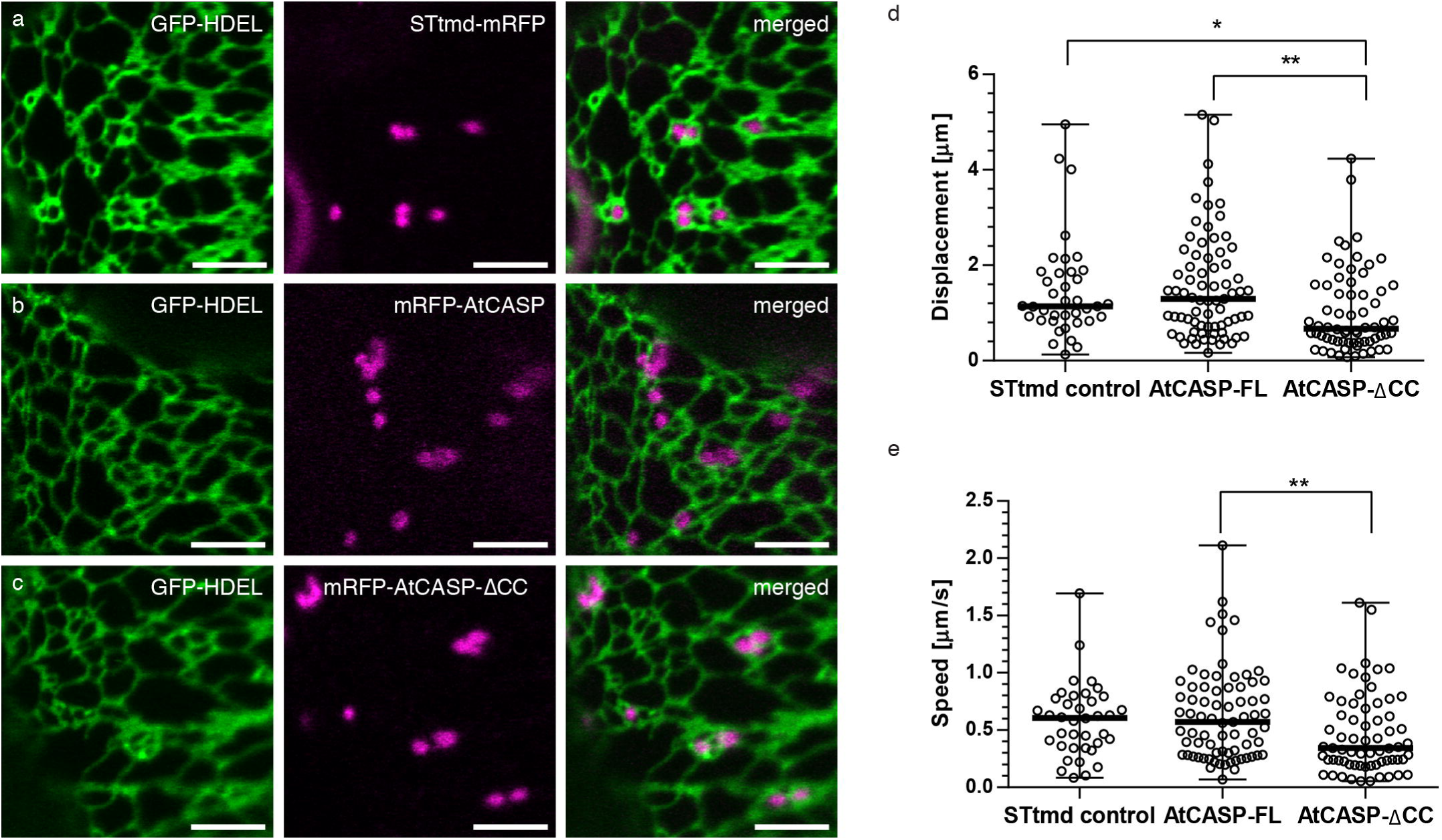
Live-cell imaging and quantitative analysis of Golgi body dynamics in AtCASP full-length and mutant *Arabidopsis thaliana* lines. a) - c) Confocal laser scanning micrographs of *Arabidopsis* cotyledonary leaf cells stably expressing the endoplasmic reticulum (ER) marker GFP-HDEL (green) and a) STtmd-mRFP, b) mRFP-AtCASP or c) mRFP-AtCASP-ACC (magenta). No obvious differences in Golgi body morphology, location or dynamics could be observed through qualitative live-cell imaging. Scale bars = 5 μm. d) - e) Quantitative analysis of d) mean speed and e) mean displacement of fluorescently labelled Golgi bodies in stable *Arabidopsis* lines expressing either the control STtmd-mRFP, mRFP-AtCASP, or mRFP-AtCASP-ΔCC. The mean speed and displacement of individual Golgi bodies were determined manually using the Fiji particle tracking plugin MtrackJ (Meijering et al. 2012). Mean speed and displacement values per cell were calculated from pooled Golgi body values (n of Golgi bodies per video ranged between 317). Statistical tests (one-way ANOVA and unpaired two-tailed student t-test) were then performed on the pooled cell values (n of cells STtmd =41, AtCASP-FL= 79, AtCASP-ΔCC = 63, see Table 1 for full summary). Scatter plots depict the mean as horizontal bar, error bars depict the SD. Asterisks represent the level of significance (^*^ p <0.05, ^**^ p= < 0.01).

**Table 1.** Numbers of *Arabidopsis* lines, cells and Golgi bodies used for analysis of velocity and displacement.

| Line | Independent lines | Individual plants per line | Repetitions per line | Total n of cells | Total n of Golgi bodies |
| --- | --- | --- | --- | --- | --- |
| STtmd-mRFP/GFP-HDEL | 1 | 5 | 3 | 41 | 332 |
| mRFP-AtCASP/GFP-HDEL | 2 | 7 and 4 | 3 and 2 | 46<br>32 | 388<br>240 |
| mRFP-AtCASP-ΔCC/GFP-HDEL | 2 | 6 and 4 | 3 and 2 | 43<br>23 | 456<br>174 |

To obtain quantitative data, movies were taken from control, full-length and mutant epidermal leaf cells and analysed using automated particle tracking software. Golgi bodies in mRFP-AtCASP lines formed clusters or chains, often just temporary in nature, with clusters dissolving after a few seconds and individual Golgi bodies continuing to move along their single trajectories (Figure 2b). Golgi body movement was therefore analysed manually using the MTrackJ plugin (Meijering *et al.,* 2012) in the ImageJ processing package Fiji (Schindelin *et al.,* 2012). MtrackJ allows the manual tracking of individual Golgi bodies frame by frame and the software was used to determine the mean speed and displacement (the straight line distance from the start point of the track to the current point measure, abbreviated here as D2S) for individual Golgi bodies in control, full-length and mutant AtCASP lines.

Mean Golgi body speed and displacement were calculated from the pooled Golgi body data (n ranging between 3-19 Golgi bodies per cell). Table 1 summarises the number of individual lines, cells and Golgi bodies that were analysed. All Golgi body values were pooled, and statistical analysis was performed on the data (one-way ANOVA, followed by an unpaired two-tailed student t-test). Scatter plots depict individual data points as well as the median Golgi body displacement and corresponding standard deviation (SD) in μm. The control lines (median = 1.14 μm, ranging from 0.13 to 4.95 μm) and full-length AtCASP lines (median = 1.24 μm, ranging from 0.16 to 5.15 μm) did not significantly differ from each other (p = 0.7256) (Figure 2d). Golgi displacement in mutant lines (median = 0.67 μm, ranging from 0.08 to 4.24 μm) was reduced significantly compared to both control (p = 0.0184) and full-length (p = 0.0020) lines. Similarly, as summarised in Fig. 2e, the mean Golgi speed did not differ significantly between control (median = 0.61 μm, ranging from 0.08 to 1.7 μm, p = 0.0740) and full-length AtCASP (median = 0.57 μm, ranging from 0.07 to 2.11 μm) lines (p = 0.5466), but was significantly decreased in the mutant (median = 0.34 μm, ranging from 0.05 to 1.61 μm) compared full-length lines (p = 0.0088).

### Optical trapping reveals that AtCASP is involved in tethering events at the Golgi-ER interface

Since Golgi movement parameters in the AtCASP lines were significantly different to the control line, optical tweezers were used to physically probe whether these were due to effecting the interaction with the ER. We hypothesised that if AtCASP had a role in tethering Golgi bodies to the ER, any potential effects of mutant AtCASP-ΔCC overexpression would become apparent upon manipulation of Golgi bodies with optical tweezers *in planta.* The underlying physical principle of optical trapping is that a highly focused laser beam is able to trap particles if they are a certain size (approx. 1 μm), and their refractive index is different to that of their environment (Neuman and Block, 2004). Golgi bodies fulfil these requirements, as their size is around 1 μm in diameter and due to their condensed stack structure their refractive index differs from the surrounding cytoplasm. In contrast, it has not been possible experimentally to trap ER membranes (Sparkes *et al.,* 2009b).

Optical trapping was performed in *Arabidopsis* cotyledonary leaf epidermal cells of four to five day old seedlings (before start of growth stage 1 and emergence of rosette leaves,Boyes *et al.,* 2001), of mRFP-AtCASP-ΔCC/GFP-HDEL (n of cells = 10, n of Golgi bodies = 45) and ST-mRFP/GFP-HDEL control lines (n of cells = 13, n of Golgi bodies = 53, Table 2). A new Golgi body was randomly chosen for every new trapping event. Leaf samples were treated with the actin-depolymerising drug Latrunculin B before trapping, to inhibit actin-based Golgi movement. Any subsequent movement was therefore due to the physical micromanipulation of the trapped Golgi body, as the ER cannot be trapped (Sparkes *et al.,* 2009b). In STtmd-control cells, a trapping laser output of 70 mW was required to trap Golgi bodies, and no trapping was possible with outputs below this. In contrast, Golgi bodies in the AtCASP-ΔCC mutant line could easily be trapped with the laser power set to 30 mW (Table 2). From the total number of Golgi bodies trapped in the mutant line, 17 Golgi bodies moved just a few μm over the ER and then came to a halt, whereas the rest could be moved over a longer distance across the cell. ER remodeling along the tracks of trapped Golgi bodies occurred only in 15 instances in the mutant (28%), compared to 41 instances in the control trapping events (91%). Sixteen Golgi bodies in the mutant line detached from GFP-HDEL-labelled tubules during the trapping event, and 11 of them re-attached to the ER as they were being moved. For the remaining trapping events it was not possible to determine whether ER reattachment took place.

**Table 2.**
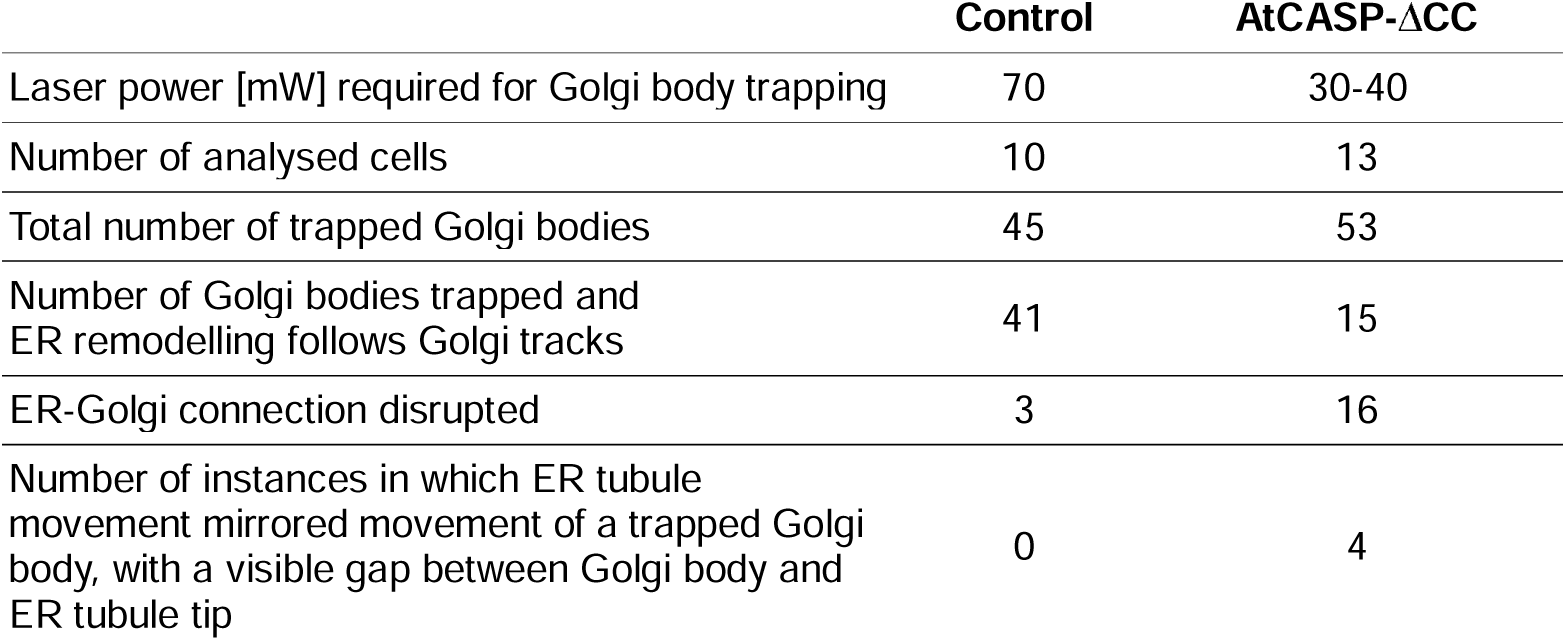
The effects of optical trapping in control and AtCASP mutant *Arabidopsis* leaf epidermal cells.

|  | Control | AtCASP-ΔCC |
| --- | --- | --- |
| Laser power [mW] required for Golgi body trapping | 70 | 30-40 |
| Number of analysed cells | 10 | 13 |
| Total number of trapped Golgi bodies | 45 | 53 |
| Number of Golgi bodies trapped and ER remodelling follows Golgi tracks | 41 | 15 |
| ER-Golgi connection disrupted | 3 | 16 |
| Number of instances in which ER tubule movement mirrored movement of a trapped Golgi body, with a visible gap between Golgi body and ER tubule tip | 0 | 4 |

Figure 3a depicts movie frames from an optical trapping event in mutant AtCASP-ΔCC cells (see Suppl. Movie 1). Turning the trapping laser on resulted in movement of a whole group of Golgi bodies over a short distance (at time point 7.8 s). A single Golgi body remained trapped (arrowhead), lost its ER tubule association and then moved freely through the cell, until connection was re-established near an ER tubule upon release of the optical trap (Fig. 3b, asterisk).

**Figure 3.**
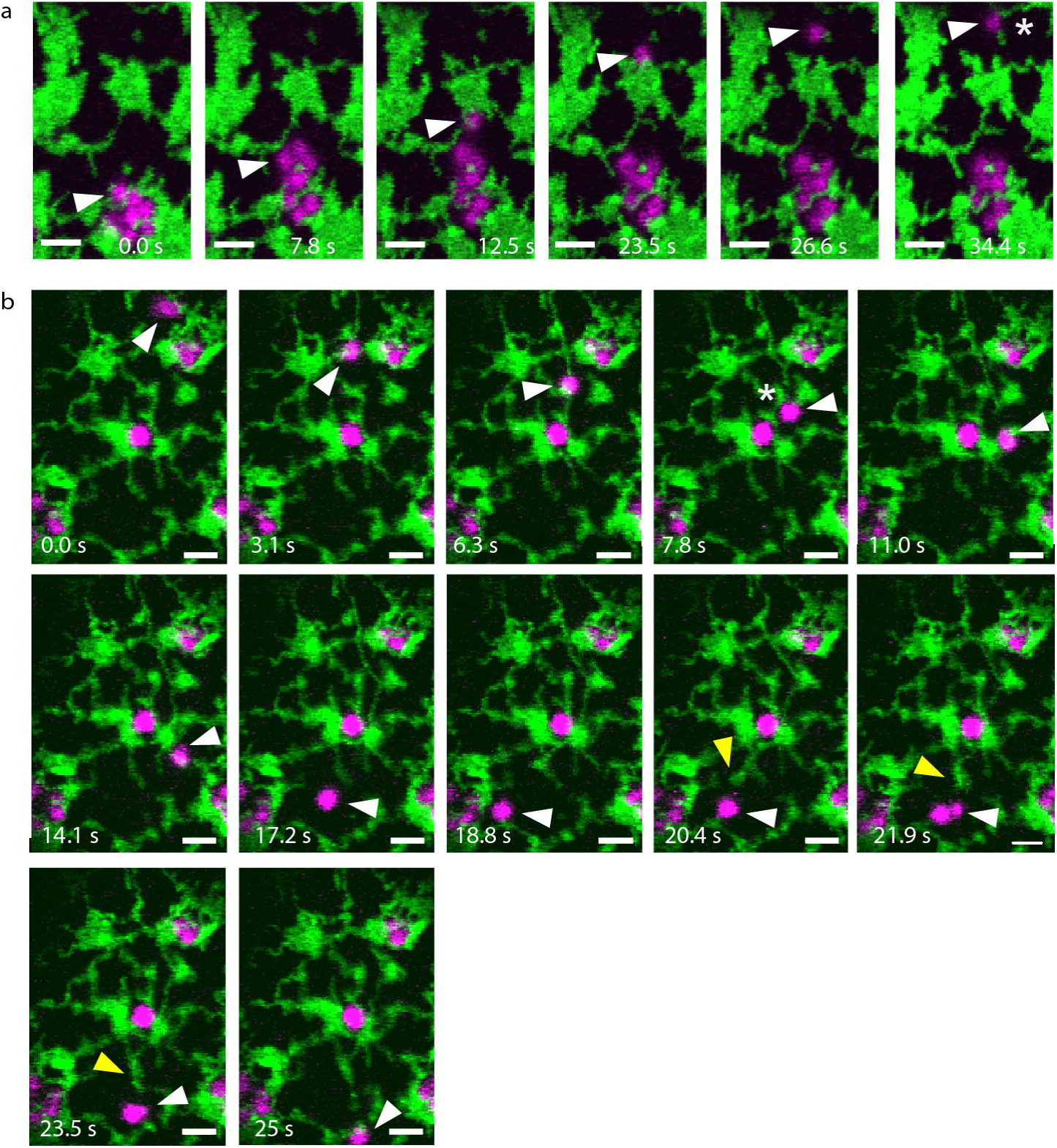
Disruption of the ER-Golgi connection in mutant AtCASP-ΔCC cells. Confocal images showing still images of a time series over 34.4 seconds during optical trapping of Golgi bodies in transgenic *Arabidopsis* cotyledonary leaf epidermal cells. Plants were expressing mRFP-AtCASP-ΔCC (magenta) and the ER marker GFP-HDEL (green). Arrowheads point to optically trapped Golgi bodies. Scale bars = 2 μm. (a) Several Golgi bodies moved with the trap across a short distance. A single Golgi body remained in the trap and moved through the cell detached from the ER. (b) A Golgi body was trapped and the ER-Golgi connection was disrupted at time point 7.8 s (asterisk). The ER tubule followed the Golgi body with a gap. At time point 20.4 s, a second ER tubule mirrored Golgi body movement with a similar gap (arrowhead).

Surprisingly, in a few instances GFP-HDEL tubules appeared to follow Golgi bodies with a significant gap after the connection had been disrupted, as shown in Figure 3b (and Suppl. Movie 2). Movement of two Golgi bodies that were trapped simultanously (Fig. 3b, arrowhead) initially resulted in ER remodeling, until the connection broke (time point 7.8 s, asterisk). The ER tubule mirrored Golgi body movement with a delay (time points 11 s to 16.8 s). From time point 20.4 s onwards, a second ER tubule mirrored Golgi body movement (yellow arrowhead), appearing to attempt attachment to the trapped Golgi body.

Interestingly, the optical trapping data mirrored the observation made during the tracking of Golgi bodies in cells expressing full-length mRPF-AtCASP, in which Golgi bodies appeared to be ‘sticky’ and formed clusters or chains. In 64% of all trapping events performed in full-length lines, two or more Golgi bodies were trapped and moved together, in contrast to 35% in STtmd-mRFP and 47% in AtCASP-ΔCC lines (Fig. 4a, at similar optical trapping force).

**Figure 4.**
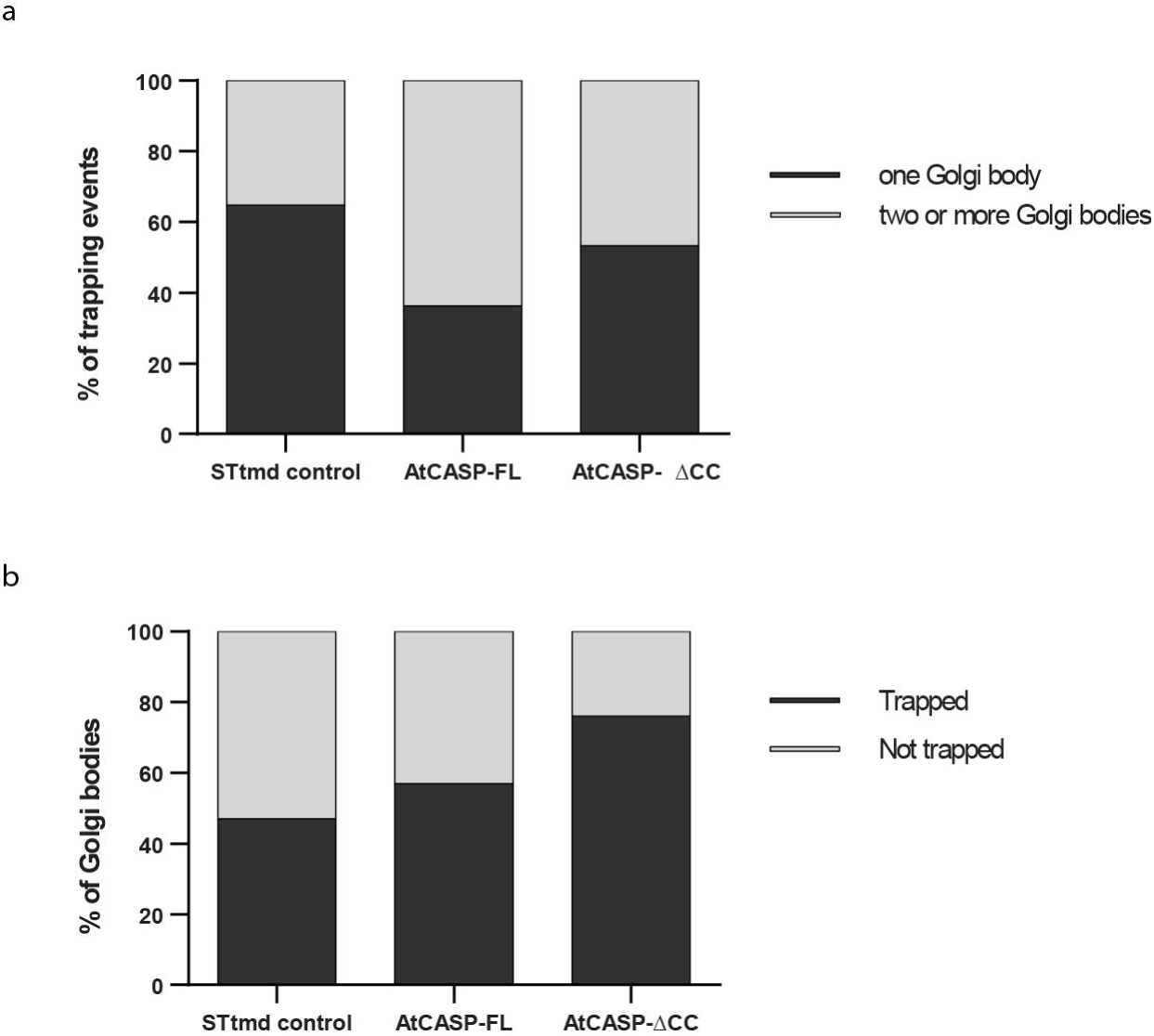
Comparing the ability to trap Golgi bodies in STtmd-mRFP control, full-length mRFP-AtCASP and mutant mRFP-AtCASP-ΔCC lines. a) Two or more Golgi bodies were captured in 64% of trapping events in full-length AtCASP lines, compared to just 35% in control, and 47% in AtCASP-ΔCC lines. Expression of full-length mRFP-AtCASP appears to make Golgi bodies ‘stickier’.
b) Average numbers of three experiments (total n=300) of trapping control, full-length and mutant AtCASP Golgi bodies in *Arabidopsis* cotyledons using a TIRF-Tweezer system. Compared to 46% trapped Golgi bodies control cells and 57% trapped Golgi in mRFP-AtCASP cells, 76% of Golgi bodies expressing the truncation could be trapped. The STtmd control and AtCASP full-length line did not significantly differ from each other (Chi-square test, p = 0.321), but the AtCASP-ΔCC line differed significantly from the control (p=1.065x10−8) and the full-length line (p=2.091×10−6)

To test the degree of attachment in more detail, we used a TIRF-based optical trapping system and captured Golgi bodies in control, full-length and mutant AtCASP *Arabidopsis* cotyledons at similar trapping force range. For this experiment, 100 Golgi bodies in three different leaves for each line (total n = 300) were randomly selected and scored as to whether they could be manually trapped and moved, or not (Figure 4c). In STtmd-mRFP cells, just 47% of Golgi bodies could be trapped, in cells expressing mRFP-AtCASP the ability to trap Golgi bodies increased slightly to 57%. In contrast, in cells expressing mRFP-AtCASP-ΔCC, we were able to trap 76% of Golgi bodies. Statistical analysis (oneway ANOVA and unpaired two-tailed student t-test) showed that there was no significant difference between the control and full-length AtCASP (p=0.321), but mRFP-AtCASP-ACC differed significantly from the control (p = 1.065×10^−8^) and full-length mRFP-AtCASP (p = 2.091×10^−6^).

Using ImageJ and the MTrackJ plugin, we mapped tracks of captured Golgi bodies in relation to the tip of the remodelling ER tubule in STtmd-mRFP control lines (five tracks in total, representative track shown in Fig. 5a), mRFP-AtCASP (six tracks, representative track shown in Fig. 5b) and mRFP-AtCASP-ΔCC lines (21 tracks, Fig. 5c and d). Arrowheads indicate trapped Golgi bodies. In control cells, Golgi body and ER tracks overlaid almost perfectly with each other during micromanipulation (n of cells = 7, Fig. 5a and e, Suppl. Movie 3). Looking at tracks from cells expressing full length mRFP-AtCASP (n = 6), we found that Golgi and ER tracks mirrored each other as they did in the control, but the connection was more easily disrupted (Fig. 5 b and f, Suppl. Movie 4) compared to the control. In cells expressing mRFP-AtCASP-ΔCC (n = 6), the instability of the ER-Golgi connection was reflected in a non-uniform range of track patterns. For example, as shown in Fig. 5 c and h (Suppl. Movie 5), an ER tubule initially followed a trapped Golgi body (arrowhead) on the same trajectory. The connection was then lost (asterisk), but the ER continued to mirror the Golgi body track but separated from each other by a distance ranging from 0.6-1.6 μm. In other instances, a captured Golgi body separated from the ER (arrowhead, Fig. 5 d and h), would reconnect with the ER for the track length of a few microns (asterisk) and break free again.

**Figure 5.**
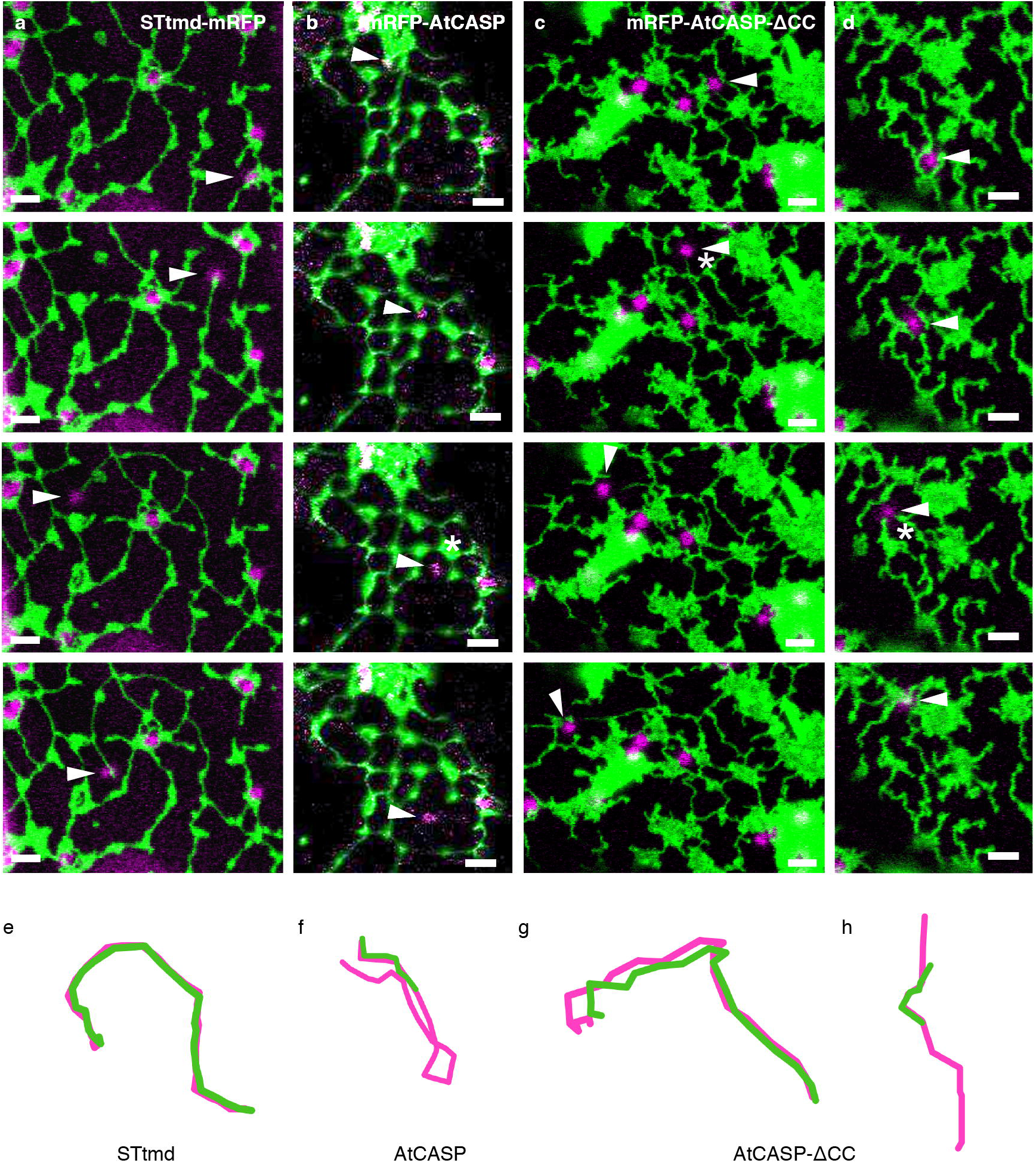
ER and Golgi body tracks differ between control and mutant lines. (a-d) Confocal images showing the effect of optically trapping individual Golgi bodies in *Arabidopsis* cotyledons expressing GFP-HDEL (shown in green) and (a) the control marker STtmd-mRFP, (b) full-length GFP-AtCASP or (c-d) truncated GFP-AtCASP-ΔCC (all shown in magenta). (e-h) Visualisation of Golgi body tracks (magenta) in relation to the ER tubule tip (green). Arrowheads indicate trapped Golgi bodies. Scale bars = 2 μm. (a) and (e) Control cell expressing STtmd-mRFP and GFP-HDEL. The Golgi-ER connection remained intact and both tracks were closely associated. (b) and (f) Cell expressing mRFP-AtCASP and GFP-HDEL. Golgi and ER remained connected only for a short time before the connection was disrupted (asterisk). (c) and (g) Cell expressing mRFP-AtCASP-ΔCC and GFP-HDEL. ER and Golgi moved together for the first part of the time series. The connection then broke apart (asterisk) and the ER followed the Golgi body with a gap. (d) and (h) Time series showing an example in which the ER-Golgi connection was disrupted immediately after trapping. A second ER tubule unsuccessfully attempted to reconnect with the Golgi body (asterisk).

The gap width between the centre of the trapped Golgi body and the ER tubule tip in the time series depicted in Fig. 5c varied throughout the optical trapping event. The distance was measured in each of the nine frames in the movie. Values ranged between 0.62 μm at the beginning to 1.33 μm at the end, with a mean width of 1.14 μm.

We assessed the stability of the ER-Golgi connection per individual trapping time series in control, full-length and mutant AtCASP lines (Fig. 6a) by calculating the ratio of frames with an intact ER-Golgi connection versus the total frame number, working on the assumption that trap movement was reasonably consistent over the short distances Golgi bodies were moved. Thus, a ratio of 1 means that ER remodelling took place throughout the whole trapping event, whereas a ratio of 0.5 indicates that the trapped Golgi body was detached from the ER for half of the time series. In control cells, 95% of trapping events showed a ratio of 1 (n=17), which reflects a stable ER-Golgi connection. In contrast, just 55% of trapped Golgi bodies in cells expressing mRFP-AtCASP (n=11), and 40% in mRFP-AtCASP-ΔCC cells (n=15) retained a permanent connection to the ER throughout the trapping event. The difference in length of disruption between the control and the full-length (p = 0.0031) or mutant AtCASP (p = 0.007) lines was significant, as determined by one-way ANOVA and unpaired two-tailed student-t test.

In control cells, only 10% of trapped Golgi bodies lost their ER connection, and if they did, it occurred just once (Fig. 6b). In full-length AtCASP expressing cells, 60% of trapped Golgi bodies detached from the ER once, which was significantly higher than control cells (one-way ANOVA and unpaired two-tailed student t-test, p = 0.0047). The ER-Golgi connection was most unstable in mRFP-AtCASP-ΔCC lines. In these, 40% of trapped Golgi bodies detached and reattached to the ER more than once during one trapping event, up to five times in one instance. This was significantly different to the control (p = 0.012), but not significantly different to full-length AtCASP (p = 0.356).

**Figure 6.**
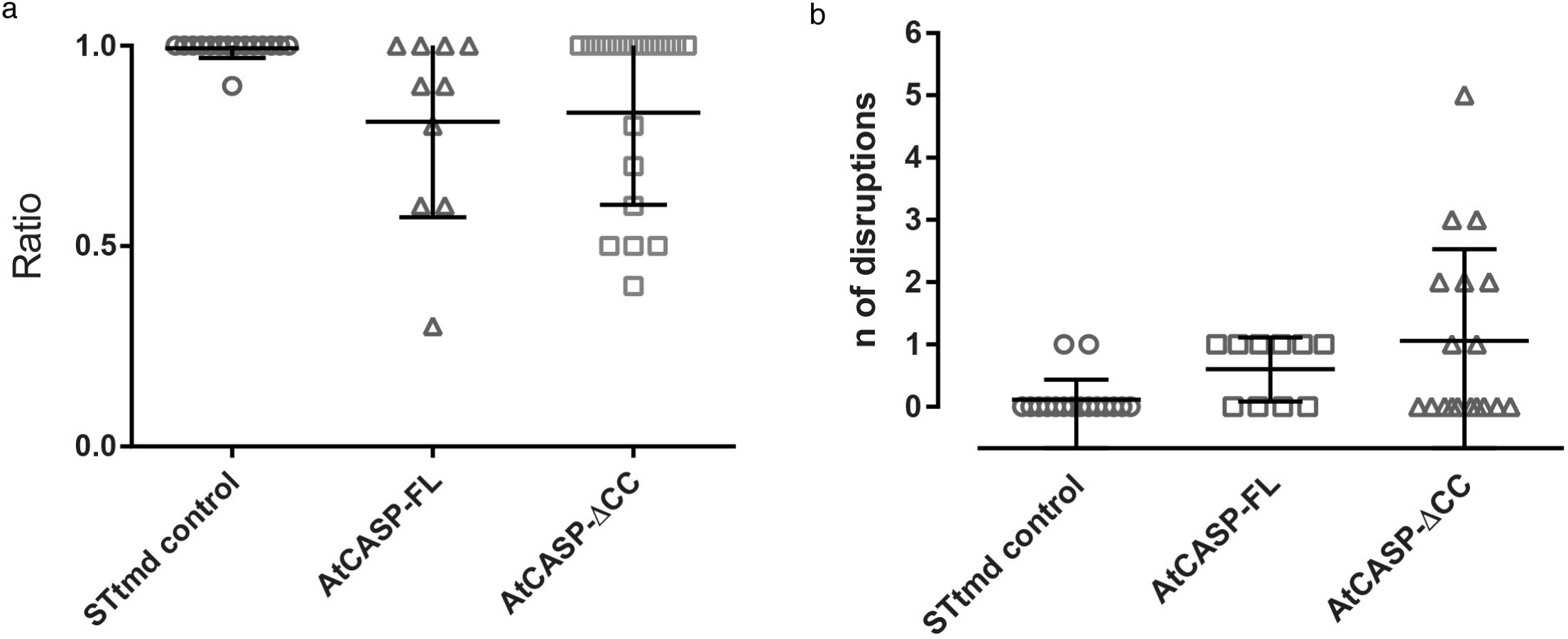
Semi-quantitative analysis of Golgi body trapping in control and AtCASP full-length and mutant expressing *Arabidopsis* lines. Assessing the stability of the connection between individual Golgi bodies and the ER in *Arabidopsis* cotyledonary leaf epidermal cells expressing STtmd-mRFP/GFP-HDEL (control, n=17), full-length mRFP-AtCASP/GFP-HDEL (n=11) or truncated mRFP-AtCASP-ΔCC/GFP-HDEL (n=15). Errors bars depict means and standard deviations. a) Scatterplot displaying the ratio of number of frames per trapping event with an intact ER-Golgi connection versus the number of total frames. A ratio of 1 indicates an intact connection over the whole duration of the time series. The smaller the ratio, the longer the connection was disrupted during a time series. The ER-Golgi connection was disrupted significantly longer in cells expressing mRFP-AtCASP (p = 0.0031) or mRFP-AtCASP-ACC (p = 0.007), compared to control cells. Full-length and mutant AtCASP lines did not differ significantly (p= 0.75). b) Scatterplot showing the number times that the ER-Golgi connection was disrupted per individual trapping event. In almost all of the trapping events in control cells, the connection remained intact. Its instability (symbolised by repeated detachments and reattachments of the trapped Golgi body with the ER) increased significantly in mRFP-AtCASP cells (p = 0.0047) and mRFP-AtCASP-ΔCC cells (p = 0.012). No significant difference was observed between full-length and mutant AtCASP (p = 0.356).

## Discussion

The advent of fluorescent protein technology permitted for the first time the observation of the dynamics of plant Golgi stacks in living plant cells (Boevink *et al.,* 1998). The movement of individual Golgi bodies over the cortical actin network whilst being somehow attached to the ER was termed “stacks on tracks”. Subsequently it was shown that transport of cargo between the ER was not dependent on the cytoskeleton (Brandizzi *et al.,* 2002), that this association encompassed the protein components of the ER exit site (da Silva *et al.,* 2004), and that the ER membrane itself was motile as well as Golgi bodies (Runions *et al.,* 2006; Sparkes *et al.,* 2009b). However, it took the application of optical trapping to conclusively demonstrate that the organelle-to-organelle adhesion at the ER-Golgi body interface was sufficiently strong to permit remodelling of the tubular ER network simply by moving Golgi bodies around in the cortex of leaf epidermal cells (Sparkes *et al.,* 2009b). Here we show that overexpression of truncated AtCASP (Latijnhouwers *et al.,* 2007; Renna *et al.,* 2005) interferes with ER-Golgi physical interaction, showing that 1) ER and Golgi bodies are tethered rather than being connections maintained through membranous extensions and 2) the ER-Golgi interface may be organised by tethering proteins, the disturbance of one, AtCASP, resulting in an alteration of Golgi movement and trapping properties.

### AtCASP functions as a tether between the ER and Golgi stack

It could be predicted that if a protein is involved in tethering the Golgi stack to the ER, then the parameters describing its movement with or over the ER may change upon its disruption. Visually, this is difficult to assess from confocal time-lapse image series, other than the observed clumping of Golgi stacks in *Arabidopsis* lines expressing full-length mRFP-AtCASP. This clumping presumably occurs due to interactions between excess coiled-coil domains on the Golgi surface. Quantitative image analysis revealed both a drop in Golgi body velocity and reduction in their mean displacement in AtCASP-ΔCC expressing cells, compared to non-clumped Golgi bodies in this AtCASP mutant or in control ST-mRFP expressing cells. This could be interpreted either as interference with putative motor protein activity at the ER Golgi interface or a loosening of the tethering at the interface. If in this scenario the tether is loosened, then decoupling of the Golgi body from its ER exit site supports the contention that Golgi movement is at least in part generated via movement of the ER surface (Runions *et al.,* 2006), in which the exit site is embedded. Alternatively, the movement of the Golgi attached to the ER exit site may affect ER movement. It is still unclear to what extent the movement of ER and Golgi are dependent upon one another, co-regulated events or mutually exclusive processes (Sparkes et al. (Sparkes *et al.,* 2009a). To date there is little evidence for a Golgi-associated myosin (Avisar *et al.,* 2009; Sparkes *et al.,* 2008), other than a study on the expression of a truncated myosin, which occasionally labelled Golgi stacks (Li and Nebenfuhr, 2007). Furthermore, the differences in the AtCASP-ΔCC mutant line were observed upon actin depolymerisation during the trapping experiments. Therefore, it can be assumed that interfering with tethering may be the most likely cause of the change in Golgi body motility on expression of mutant AtCASP.

In this study, we utilised a more direct approach to probe Golgi tethering to the ER, which was carried out on two different optical trapping set-ups, confocal and TIRF-based. Our optical trapping data clearly demonstrates that interfering with the coiled-coil domain, and thus with any tethering function, affects the physical Golgi-ER connection. We were able to show that upon overexpression of the truncated AtCASP protein, the trap power required to manipulate individual Golgi stacks was greatly reduced from that required for wild type Golgi bodies marked with a different membrane construct. Presumably, truncated AtCASP out-competed the native protein in a dominant-negative fashion. We found that trapping of Golgi bodies in mutant lines was easier, and that the interface between ER and Golgi could be disturbed under experimental conditions in which actin had been depolymerised.

### AtCASP: One component of a larger tethering complex?

In control Golgi-tagged plants, upon micromanipulation of Golgi bodies, the ER track coincided almost perfectly with the Golgi track. Upon over expression of full-length fluorescently tagged AtCASP, the connection appeared to be more easily disturbed than in control cells, but the tracks of ER tips and Golgi bodies occasionally were able to mirror each other. Golgi bodies still appeared to move on actin delimited ‘tracks’, but the connection with the ER was loose. In cells expressing the deletion mutant, the disruption of the putative tether was obvious. Golgi bodies broke free from the ER more easily than in control STtmd-mRFP or full-length mRFP-AtCASP expressing cells. Track patterns were irregular and did not mirror that of the ER.

The gap observed on some occasions whilst being a micron plus between Golgi and ER, showed the Golgi and ER following the same trajectory, suggests that AtCASP is not solely responsible for tethering at the ER-Golgi interface, but might be part of a more substantial tethering complex. Other components of such a complex might include other *cis*-Golgi located golgins such as the plant homologue of the well-characterised tether Atp115 (MAG4,Kang and Staehelin, 2008; Lerich *et al.,* 2012; Takahashi *et al.,* 2010), or even the recently identified AtSec16/MAIGO5 (Takagi *et al.,* 2013). Gillingham and colleagues (Gillingham *et al.,* 2002)reported an indirect interaction between the yeast CASP homologue COY1 and the SNARE (Soluble N-ethylmaleimide-sensitive factor Activating protein REceptor) protein Gos1p in yeast assays, as well as a small fraction of COY1 co-precipitating with the COPII coat subunit hSec23 and Golgin-84. Another study (Malsam *et al.,* 2005) identified mammalian CASP as component of an asymmetric tethering complex, with CASP binding to Golgi membranes and interacting with Golgin84 on COPI vesicles, thus suggesting a role for CASP in retrograde transport. As many protein functions within the secretory pathway are conserved between plants and mammals, some interactions might be conserved as well, and we are currently analysing potential AtCASP binding partners.

### AtCASP as novel starting point to dissect the plant ER-Golgi interface

Previous studies suggested a role for AtCASP in Golgi biogenesis, possibly as part of a ‘platform’ that might act as base for the formation of early *cis*-Golgi structures (Ito *et al.,* 2012; Osterrieder *et al.,* 2010; Schoberer *et al.,* 2010). As immuno-labelling of GFP-AtCASP located the construct to cisternal rims of Golgi stacks (Latijnhouwers *et al.,* 2007), AtCASP appears to be anchored through its transmembrane domain to *cis*-Golgi membranes, while its coiled-coil domains (labelled by the N-terminal fluorophore) bind to yet unidentified partners at the ER-Golgi interface. Triple labelling experiments with fluorescent full-length and mutant fluorescent AtCASP versions, co-expressed with ER exit site and COPII markers, as well as *cis-* or *trans-Golgi* membrane markers such as glycosyltransferases (Schoberer and Strasser, 2011), could help to unravel the subcompartmentalisation of key players at the plant ER-Golgi interface.

The biology of the ER-Golgi interface differs between plants and mammals in a variety of aspects (Brandizzi and Barlowe, 2013). Based on this and on the results from our study, we hypothesise that AtCASP could have different or additional functions in plants compared to its animal and yeast homologues. Notably, the model of ER exit site organisation itself is still in flux. The latest model, proposed by Glick (2014), replaces the concept of a COPII-organising scaffold with that of a self-organising tethering framework, consisting of ER exit sites (transitional ER sites in yeast), early Golgi membranes and tethering factors, one of which might be CASP (Glick, 2014). Understanding the molecular make-up and mechanisms of the plant ER-Golgi interface is crucial for our understanding of how proteins pass through the secretory pathway. By identifying AtCASP as novel ER-Golgi tether, we have gained a new entry point into the dissection of the plant ER-Golgi interface.

## Conclusion

In conclusion this work indicates that leaf epidermal cell Golgi bodies are intimately associated with the ER and that the connection is most likely maintained by a tethering complex between the two organelles, thus not simply relying on membrane continuity between the ER exit site and *cis*-Golgi membranes.

## Acknowledgements

We thank Janet Evins for help with growing plants. The work was supported by a BBSRC grant (BB/J000302/1) and a Royal Society Travel grant to CH.

## Short legends for supporting information (separate files)

### Suppl. Movie 1

Confocal images of a timeseries, taken over 34.4 seconds, showing *Arabidopsis thaliana* leaf epidermal cells with endoplasmic reticulum labelled with GFP-HDEL and Golgi bodies labelled with mRFP-AtCASP-ΔCC. Initially, a whole group of Golgi bodies moved with the trap. A single Golgi body remained in the trap, lost connection to the ER tubule, and then moved freely through the cell until connection was re-established near an ER tubule. Scale bar = 2 μm.

### Suppl. Movie 2

Confocal images of a timeseries, taken over 70.4 seconds, showing *Arabidopsis thaliana* leaf epidermal cells with the endoplasmic reticulum labelled with GFP-HDEL, and Golgi bodies labelled with mRFP-AtCASP-ΔCC. The optically trapped Golgi body lost its connection to the ER, its movement being mirrored by the ER with a gap between the both. Scale bar = 2 μm.

### Suppl. Movie 3

Confocal images of a timeseries, taken over 15.24 seconds, showing *Arabidopsis thaliana* leaf epidermal cells with the endoplasmic reticulum labelled with GFP-HDEL and Golgi bodies labelled with STtmd-mRFP. The movement of the trapped Golgi body and the tip of the ER tubule overlaid almost perfectly with each other during micromanipulation. Scale bar = 2 μm.

### Suppl. Movie 4

Confocal images of a timeseries, taken over 10.40 seconds, showing *Arabidopsis thaliana* leaf epidermal cells with the endoplasmic reticulum labelled with GFP-HDEL and Golgi bodies labelled with mRFP-AtCASP. Golgi and ER tubule tracks mirrored each other as in the control, but the ER-Golgi connection was more easily disrupted. Scale bar = 2 μm.

### Suppl. Movie 5

Confocal images of a timeseries, taken over 10.27 seconds, showing *Arabidopsis thaliana* leaf epidermal cells with endoplasmic reticulum labelled with GFP-HDEL and Golgi bodies labelled with mRFP-AtCASP-ΔCC. The ER tubule initially followed the trapped Golgi body. The connection became disrupted, but the ER continued to mirror the Golgi body movement. Scale bar = 2 μm.

